# Best for the Eye, Not for the Algorithm: Anisotropy in Fitting Atomic Models in Cryo-EM

**DOI:** 10.64898/2026.07.15.738625

**Authors:** Roey Yadgar, Roy R. Lederman

## Abstract

Most atomic model refinement methods in cryo-EM fit models to the reconstructed density map and effectively treat Fourier voxels as equally reliable. However, the uncertainty in the estimation of Fourier coefficients is highly anisotropic, primarily due to the common variability in SNR in different frequency shells and the distribution of particle images across viewing directions.

First-principles arguments suggest that atomic models should be fitted to particle images rather than volumes; this strategy may be computationally demanding. We show that under certain modeling choices, fitting atomic models to *weighted* volumes is equivalent to fitting directly to particle images. Furthermore, we argue that various proxies can be used to capture this and other sources of uncertainty and distortions.

We propose that the principle can be implemented in most atomic model-fitting software with relative ease, using information readily available in existing pipelines. As a proof of concept, we extracted the necessary information from standard RELION runs and fed it into a modified version of Servalcat in which we implemented a reinterpreted version of the idea.

## 1 Introduction

Cryo-electron microscopy (cryo-EM) pipelines reconstruct three-dimensional density maps from large collections of noisy two-dimensional projection images. The reconstructed map is the canonical output of the modality: investigators inspect it on a screen, deposit it in public databases, and use it as the input to atomic modeling. Nearly every flexible-fitting and refinement method in widespread use — MD-based, normal-mode-based, real-space, or reciprocal-space — treats this map as the data against which an atomic model is scored.

The map, however, is not the data. It is a derived quantity: a Fourier-space back-projection of the underlying particle stack, with each Fourier voxel populated by a CTF- and pose-weighted sum of contributions from individual particle images. The information available about any single Fourier voxel depends on how many particles landed in orientations that observe it, on those particles’ CTFs, and on the per-shell noise level. When the pose distribution is non-uniform — as it almost always is in practice — the resulting reconstruction noise structure across Fourier space is sharply anisotropic: voxels within a single frequency shell can differ in reconstruction signal-to-noise ratio by an order of magnitude or more.

Standard refinement pipelines use reconstructed volumes that are affected by this directional bias, but they discard information about it. At best, they apply a per-shell weighting; often, they treat all voxels as equally reliable. We submit that this omission is not merely a modeling shortcut: it reflects the way cryo-EM outputs have evolved around the needs of human interpretation. Sharpening, B-factor application, and FSC-weighted filtering all aim to produce maps that render more clearly for humans, conflating signal with noise across directions and producing a smoothed, isotropized object that is easy to inspect but that has had much of its uncertainty structure obscured. The map that is best for visualization does not carry the most usable information for an algorithm.

This distinction has a precise consequence. By the information-processing inequality, no function of the map can recover information that the map construction has discarded; an algorithm that consumes the unweighted map inherits that loss. However, observe that the same particle stack that yields the reconstructed map also yields — at no additional cost — a per-voxel variance, which standard reconstruction software (e.g., RELION [12]) computes internally and exposes on request. Folding this variance into the maximum-likelihood objective used by reciprocal-space refinement restores the directional information that map post-processing removes. Moreover, the reweighted likelihood on the map is (under the assumption of known poses) mathematically equivalent to the likelihood on the original particle stack: the map, together with its per-voxel variance, is a sufficient statistic for refinement, whereas the map alone is not.

We demonstrate the consequence concretely by modifying Servalcat [11], a standard refinement tool, to accept the per-voxel variance produced by RELION’s existing external-reconstruction interface; we note that we introduce a reinterpretation of the first-principles description and algorithm above to fit Servalcat’s algorithm. On synthetic streptavidin data with a non-uniform pose distribution and low signal-to-noise ratio, the modification reduces by 15% the C*α* RMSD to ground truth, and improves the model-to-map FSC. The model that best fits the visualized map is not the model closest to the truth, and the gap between the two is exactly the directional bias and the variability of the observed signal in different frequency shells, which the standard pipeline ignores.

We note that there are various ways to interpret the insight presented here. Indeed, as we noted, in our implementation, we reinterpreted the first-principles algorithm in the context of Servalcat’s framework (which models per-frequency shell variance). We propose that the directional (and radial) information can be approximated through other proxies, such as directional FSC as inferred from half-maps and even measures of the directional distribution of images. Furthermore, we note that there are additional sources of noise or distortion that are not covered by our model, for example, the effects of pose error or heterogeneity on the maps. We propose that the core idea can be implemented in different ways in most existing software packages for fitting atomic structures for cryo-EM data.

## 2 Background

Cryo-EM single-particle analysis produces a three-dimensional density map by combining many two-dimensional projection images of a frozen macromolecular complex. Extracting an atomic model from such a map requires fitting an initial model (e.g. AlphaFold [1] prediction) to the observed density. This process is commonly done in two stages that differ in the magnitude of displacements involved and in the computational strategy they employ: *flexible fitting* (e.g. [2, 3, 4, 5, 6, 7]) and *model refinement* (e.g. [9, 10, 8, 11]). At the risk of oversimplifying a wide array of procedures and approaches, we refer to them collectively as atomic model fitting, as the principles presented in the paper are applicable in both settings.

### 2.1 A Common Limitation of Existing Methods

A shared limitation of all these approaches is that the Fourier voxels of the experimental density map are treated as equally reliable, despite the well-known variability and anisotropy in reconstruction noise. Standard refinement pipelines use single particle reconstruction algorithms (e.g. RELION [12] or cryoSPARC [13]) to produce a density map from a particle stack. This particle stack naturally contains less information in the higher frequency components (as a consequence of the Fourier Slice Theorem [14]). Moreover, some Fourier voxels within the same shell may have higher or lower reconstruction signal-to-noise ratio (SNR) than others, an effect that is especially pronounced when the particle stack contains a non-uniform pose distribution. In other words, the reconstructed map very often has an anisotropic noise structure.

The methods mentioned above do not use the particle stack information, but instead take the final reconstructed maps as input, and treat all Fourier voxels as equally reliable. In some cases, they use heuristic weighting per frequency shell: a radial weight profile that does not account for non-uniformity of the poses in the particle stack.

We propose a simple modification to the standard ML objective that incorporates the per-voxel noise variance, which can be computed from the particle stack without any additional data collection or processing steps. This modification leads to a more accurate structural fit, especially in low-SNR regimes where the noise is highly anisotropic across Fourier space.

### 2.2 The Maximum-Likelihood Target

We consider a simplified ML formulation of fitting atomic models to cryo-EM data; for clarity, we assume that the particle poses are known.

The imaging model is:

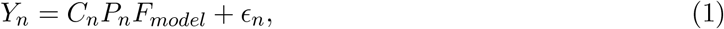

where *Y*_*n*_ is the Fourier transform of the *n*-th particle image, *C*_*n*_ is its contrast transfer function (a diagonal matrix in Fourier space), *P*_*n*_ is the slicing operator that extracts the 2D Fourier slice of the 3D volume at the *n*-th particle’s orientation, *F*_*model*_ is the Fourier transform of the model’s simulated density (typically approximated as a sum of Gaussians at the atomic positions), and *ϵ*_*n*_ is the noise in the *n*-th image, with isotropic noise within each frequency shell: 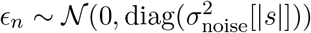.

The exact form of the likelihood under this imaging model involves operator-weighted terms. Under the nearest-neighbor discretization of the projection operator—a standard approximation in cryo-EM that makes the reconstruction problem diagonal in Fourier space—the reconstructed Fourier voxel at frequency index *k* is:

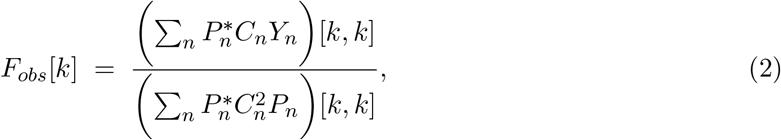

where *F*_*obs*_ is the observed reconstructed volume, and [*k, k*] denotes the *k*-th diagonal entry of the corresponding matrix. Combining (1) with (2) (see Appendix A), the reconstructed Fourier coefficient follows:

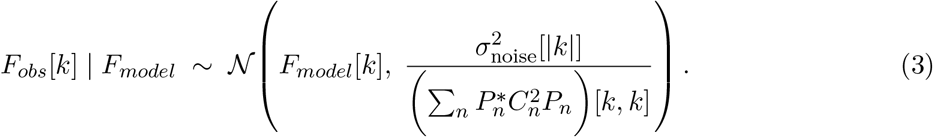

The per-voxel variance in (3) depends on the particle pose distribution, the CTFs, and the per-shell noise level in the particle images, and is therefore highly anisotropic across Fourier space. Furthermore, when the pose distribution has strong preferred orientations, voxels within the same frequency shell can differ substantially in reconstruction SNR.

The negative log-likelihood of *F*_*obs*_ given *F*_*model*_ is therefore

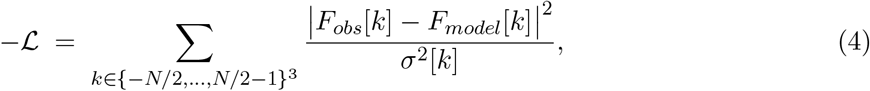

where 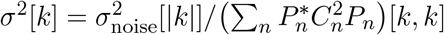.

There is a variety of expressions used in atomic fitting algorithms in practice. Many methods use the unweighted *L*^2^ objective ∥*F*_*obs*_ − *F*_*model*_∥^2^, which corresponds to the special case *σ*^2^[*k*] = const. Some apply a per-shell radial weight that captures the overall decay of signal with frequency but does not account for anisotropy within a shell, and others use the cross-correlation target 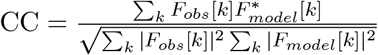 We argue that using the negative log-likelihood with the per-voxel weights derived from (3) is the principled choice, and that using constant or per-shell weights is suboptimal when the pose distribution is non-uniform.

A further consequence of (3) is that minimizing (4) with these per-voxel weights is equivalent to minimizing the negative log-likelihood of the full particle stack:

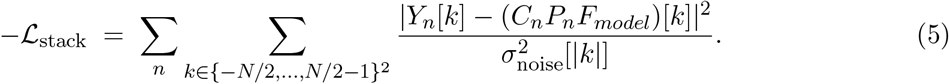

That is, the reconstructed volume together with its per-voxel variance constitutes a sufficient statistic for model fitting: no information about *F*_*model*_ is lost by working with the volume and appropriate weights rather than directly with the particle images (under known poses and the nearest-neighbor approximation).

We note, however, that the model above still makes simplifying assumptions that do not fully capture real data. Effects such as the use of real-space masking, structural heterogeneity in the data, particle pose errors and uncertainty, and model incompleteness are not reflected in 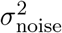 alone.

## 3 Implementation in Servalcat

We illustrate the concept of using weights derived from the particle stack by modifying Serval-cat [11], a reciprocal-space ML refinement software for cryo-EM structures. Servalcat uses the maximum-likelihood target, but introduces a per-shell scale factor *D*_*b*_ to account for the overall signal decay with frequency, and a per-shell variance *S*_*b*_ to model the noise level. This formulation is different from our theoretical model above, and therefore we reinterpret the idea in the context of Servalcat’s formulation and algorithm, where the negative log-likelihood is

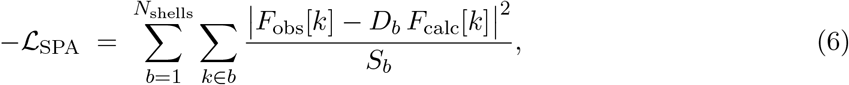

where in each refinement iteration, Servalcat updates *D*_*b*_ and *S*_*b*_ by:

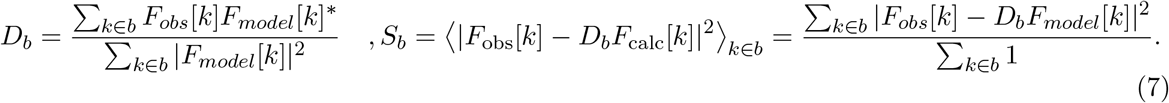

In practice, the residuals |*F*_obs_[*k*] − *D*_*b*_*F*_calc_[*k*]|^2^ contain contributions not only from measurement noise *σ* [*k*] but also from model incompleteness and coordinate error, which are not captured by the reconstruction weights.

Following the S`ERVALCAT` framework, we estimate this model-error contribution per shell and incorporate it into an *effective* per-reflection variance. Specifically, after updating *D*_*b*_ via the weighted least-squares estimator

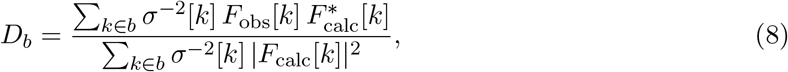

we compute the per-shell mean residual variance as in (7), and define the per-shell model-error variance as

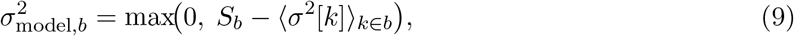

which is the excess residual variance not accounted for by the input per-reflection noise. The effective variance then replaces *S*_*b*_ in the likelihood target (6).

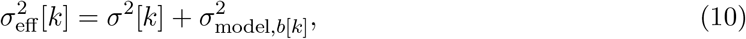

where *b*[*k*] denotes the shell containing reflection *k*. By construction, 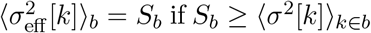(i.e. excess residual not accounted by *σ*^2^[*k*] is present), so the shell-average effective variance agrees with the standard Servalcat estimate; however, the within-shell variation is anisotropic, reflecting the directional structure of the reconstruction noise *σ*^2^[*k*].

### 3.1 CLI Integration

We added two new flags to `servalcat refine_spa_norefmac` in our modified version of Servalcat (which can be found in this fork https://github.com/RoeyYadgar/servalcat-weighted-refinement):

~~~
--relion_model_star <model.star>
--relion_weight_mrc <external_reconstruct_weight.mrc>
~~~

where `model.star` is the model file outputted by a `relion_refine` run (which contains the estimated particle image noise that RELION estimates during the refinement process), and `external_reconstruct_weight.mrc` is generated by `relion_refine` when passing the `--external_reconstruct` flag.

## 4 Validation on Simulated Data

### 4.1 Experimental Setup

To isolate the effect of the weighting scheme from other variables we performed a fully controlled simulation experiment. A particle stack of 5000 images was generated from the ground-truth atomic model of entry 7DY0 [15], with pixel size 0.8 Å, box size 110 × 110 pixels, a CTF with defocus ranging from 1.0 to 2.5 *µm*, and signal-to-noise (SNR) ranging from 10^−3^ up to 1. We evaluated the experiment in both the uniform particle poses setting, and a highly non-uniform pose distribution generated as a mixture of a von Mises-Fisher distribution (*κ* = 10) and a uniform distribution, weighted 0.95 and 0.05 respectively.

We ran the weighted modification and standard Servalcat, starting from a shaken copy of the ground-truth model, produced by adding independent Gaussian perturbations with *σ* = 0.5 Å to each atomic coordinate. Twenty iterations were run for each method with all other settings identical.

### 4.2 Results

Figure 1 shows that the weighted modification to Servalcat is able to improve the RMSD and FSC of the refined model in the settings of non-uniform poses. Additionally, the per-residue C*α* deviation histograms (Figure 2a) confirm that the improvement is distributed broadly across the chain rather than being localized to a few flexible loops, and the correlation against the ground truth model is improved the high-resolution frequency shells (Figure 2b).

**Figure 1:**
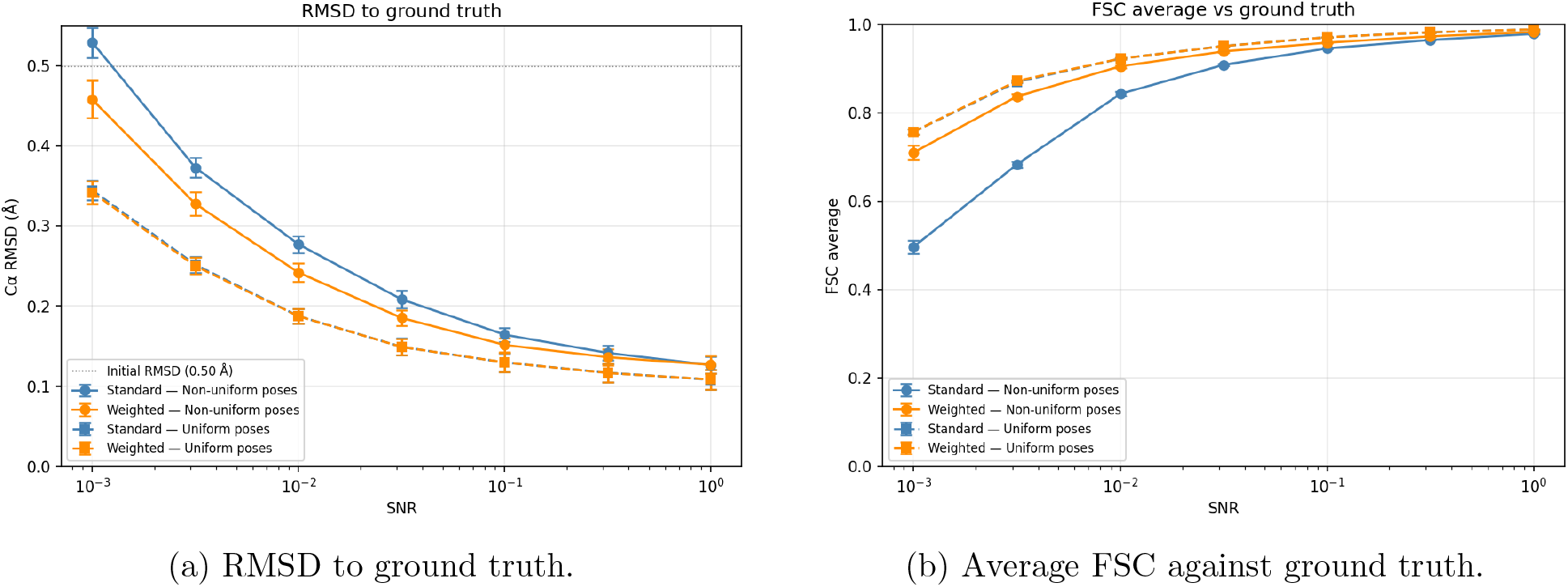
Standard vs. weighted Servalcat across SNR levels and pose distribution settings. The weighted method matches standard performance under uniform poses and improves RMSD and FSC under non-uniform poses.

**Figure 2:**
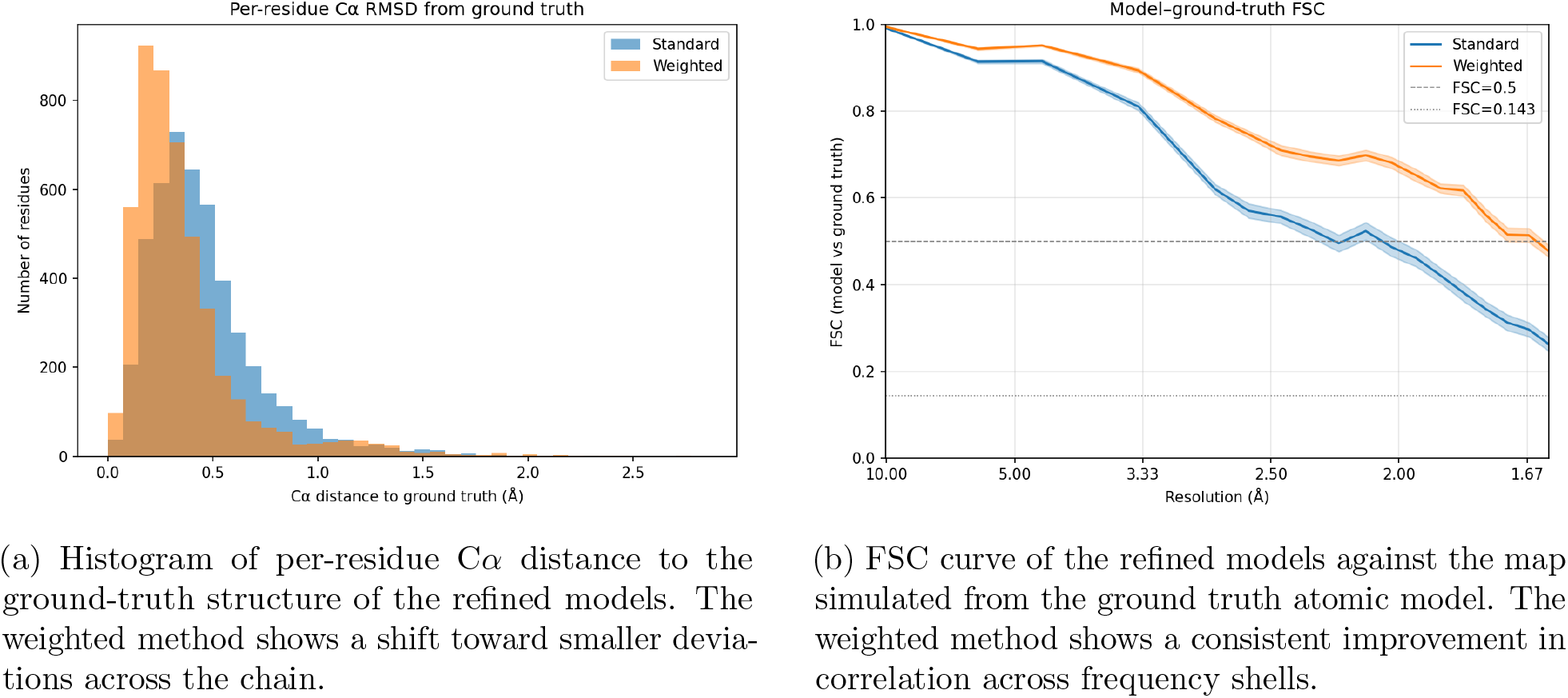
Standard vs. weighted Servalcat at *SNR* = 10^−3^ with highly non-uniform particle poses.

## 5 Conclusions

Standard refinement pipelines discard the information about the anisotropic noise structure of the reconstructed map. We have shown that this information can be folded back into the ML objective as a per-reflection variance, and demonstrated a concrete implementation within Servalcat. The approach requires no extra data-collection or processing steps beyond what RELION already produces, and numerical experiments confirm its effectiveness. While we have only implemented this approach in Servalcat, the underlying principle is general and can be applied to other refinement frameworks. Different implementations are likely to require reinterpretation of the underlying idea, as exemplified in the variation we designed to fit the Servalcat algorithm. Different implementations can also use different sources to estimate the missing information or missing uncertainty. For example, one could use half-maps to produce similar anisotropic estimates.

## Acknowledgments

We are deeply grateful to the developers of Servalcat for making their software open source; their work was instrumental in allowing us to develop and integrate our weighted refinement approach into a forked version of their repository. We also thank the Yale Center for Research Computing (YCRC) for providing computing resources and support. The work was supported by the Alfred P. Sloan Foundation (FG-2023-20853), and the Simons Foundation (1288155). Research reported in this publication was supported by the National Institute of General Medical Sciences of the National Institutes of Health under Award Number R35GM157226. The content is solely the responsibility of the authors and does not necessarily represent the official views of the National Institutes of Health.

## Appendix A Technical derivations

### Lemma 1.

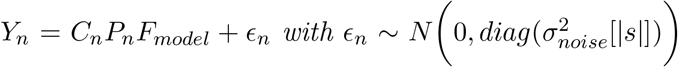 *and assume P*_*n*_ *(which contain the particle image poses) are known and fixed, then if:*

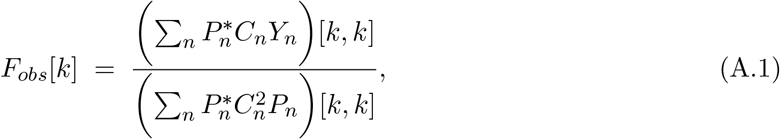

*as in* (2) *then:*

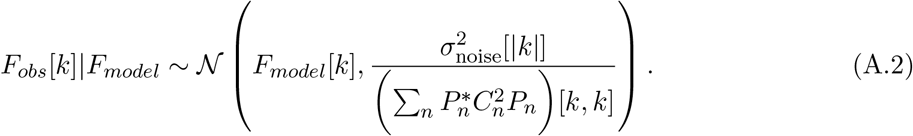

*Proof*. Since *F*_*obs*_ is linear with respect to the images *Y*_*n*_ we have:

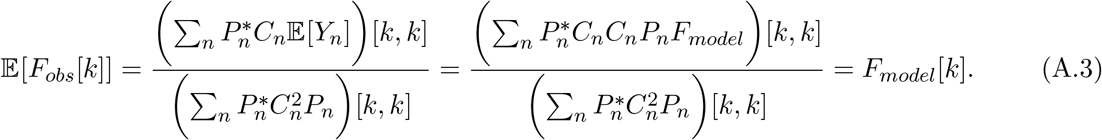

Furthermore, assuming independence of the noise *ϵ*_*n*_ in the particle images:

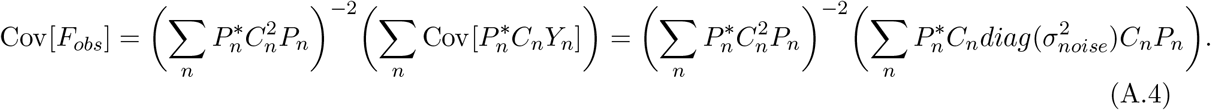

But since both *diag* 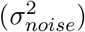 and *C*_*n*_ are diagonal and radially symmetric, we have that 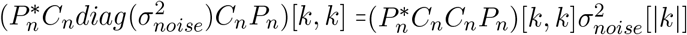,so:

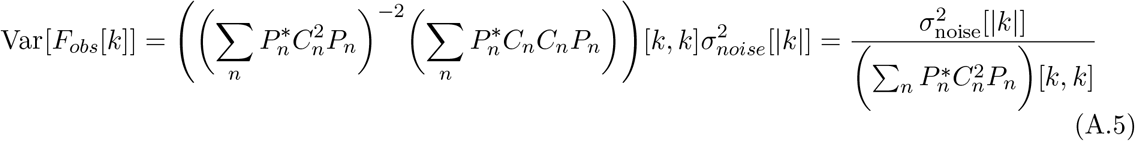

## References

[1] J. Jumper et al., Highly accurate protein structure prediction with AlphaFold, Nature 596, 583–589 (2021).

[2] F. Tama, O. Miyashita, C. L. Brooks III, Normal mode based flexible fitting of high-resolution structure into low-resolution experimental data from cryo-EM, J. Struct. Biol. 147, 315–326 (2004).

[3] L. G. Trabuco, E. Villa, K. Mitra, J. Frank, K. Schulten, Flexible fitting of atomic structures into electron microscopy maps using molecular dynamics, Structure 16, 673–683 (2008).

[4] M. Topf et al., Protein structure fitting and refinement guided by cryo-EM density, Structure 16, 295–307 (2008).

[5] J. R. López-Blanco, P. Chacón, iMODFIT: Efficient and robust flexible fitting based on vibrational analysis in internal coordinates, J. Struct. Biol. 184, 261–270 (2013).

[6] M. Igaev, C. Kutzner, L. V. Bock, A. C. Vaiana, H. Grubmüller, Automated cryo-EM structure refinement using correlation-driven molecular dynamics, eLife 8, e43542 (2019).

[7] D. N. Kim et al., cryo fit: Democratization of flexible fitting for cryo-EM, J. Struct. Biol. 208, 1–6 (2019).

[8] P. V. Afonine et al., Real-space refinement in Phenix for cryo-EM and crystallography, Acta Cryst. D 74, 531–544 (2018).

[9] Z. Wang, G. F. Schröder, Real-space refinement with DireX: From global fitting to side-chain improvements, Biopolymers 97, 687–697 (2012).

[10] A. Brown et al., Tools for macromolecular model building and refinement into electron cryomicroscopy reconstructions, Acta Cryst. D 71, 136–153 (2015).

[11] K. Yamashita, C. S. Palmer, T. Burnley, G. N. Murshudov, Cryo-EM single particle structure refinement and map calculation using Servalcat, Acta Cryst. D 77, 1282–1291 (2021).

[12] S. H. W. Scheres, RELION: Implementation of a Bayesian approach to cryo-EM structure determination, J. Struct. Biol. 180, 519–530 (2012).

[13] A. Punjani, J. L. Rubinstein, D. J. Fleet, M. A. Brubaker, cryoSPARC: algorithms for rapid unsupervised cryo-EM structure determination, Nat. Methods 14, 290–296 (2017).

[14] R. A. Crowther, L. A. Amos, J. T. Finch, D. J. DeRosier, A. Klug, Three dimensional reconstructions of spherical viruses by Fourier synthesis from electron micrographs, Nature 226, 421–425 (1970).

[15] M. Hiraizumi, K. Yamashita, T. Nishizawa, M. Kikkawa, O. Nureki, 1.93 Å cryo-EM structure of streptavidin, PDB entry 7DY0 (2021).

